# FLT3 signaling inhibition preserves opioid analgesia while abrogating tolerance and hyperalgesia

**DOI:** 10.1101/2023.03.16.532971

**Authors:** Antoine Jouvenel, Adrien Tassou, Maxime Thouaye, Jérôme Ruel, Myriam Antri, Jean-Philippe Leyris, Aurore Paquet, Sylvie Mallié, Chamroeum Sar, Lucie Diouloufet, Corinne Sonrier, François Daubeuf, Juliette Bertin, Stacy Alves, Stéphanie Ventéo, Nelly Frossard, Patrick Carroll, Ilana Mechaly, Didier Rognan, Pierre Sokoloff, Radhouane Dallel, Patrick Delmas, Jean Valmier, Cyril Rivat

## Abstract

Opioid analgesia is counteracted on chronic use by tolerance and hyperalgesia inducing dose escalation and life-threatening overdoses. Mu opiate receptors (MOR) expressed in primary sensory neurons were recently found to control tolerance and hyperalgesia, but the underlying mechanisms remained elusive. Here we show that genetic inactivation of *fms*-like tyrosine kinase receptor 3 (FLT3) receptor in sensory neurons abrogates morphine tolerance and hyperalgesia by preventing MOR-induced hyperactivation of the cAMP signaling pathway and subsequent excitatory adaptive processes. Moreover, the specific FLT3 inhibitor BDT001 potentiates morphine analgesia in acute and chronic pain models, without aggravating morphine adverse effects, and reverses tolerance and hyperalgesia once installed. Thus, FLT3 appears as a key regulator of the MOR signaling pathway and its pharmacological blockade shows promise to enhance chronic opioid analgesic efficacy.

---

Opioid analgesics remain the most effective drugs for managing moderate to severe pain, but while their efficacy is undisputed in acute and cancer pain, their long-term use for treating other chronic pain disorders has met increasing scrutiny. Chronic opioid treatment is limited by insufficient long-term efficacy (*1*), development of analgesic tolerance and paradoxical hyperalgesia (*2–4*), which lead to dose escalation, addiction and life-threatening respiratory depression (*5*). Adaptive processes responsible for the progressive decline of opioid clinical effectiveness are still inadequately managed by current drugs, indicating an urgent requirement of therapeutic strategies for alleviating associated tolerance and hyperalgesia, thus enabling improved chronic opioid analgesia.

MOR is widely distributed in neural circuits for pain and is considered as the main mediator of morphine analgesia (*2*). However, MOR expressed in dorsal root ganglion (DRG) neurons appears to be critical in the development of morphine-induced tolerance (MIT) and hyperalgesia (MIH). Indeed, selective MOR deletion in DRG neurons abrogates MIT and MIH, whereas central analgesia is maintained (*6–8*). These data indicate that MIT and MIH can be dissociated from morphine analgesia and thus compel a better understanding of how chronic opioids cause cellular plasticity and functional alteration in DRG neurons.

Neuronal pronociceptive adaptations involved in the onset of MIT and MIH show similar molecular mechanisms to those implicated in the development of peripheral neuropathic pain (PNP) (*9–12*), which is initiated and maintained by FLT3 expressed in primary sensory neurons (*13*). FLT3 activation induces maladaptive excitatory responses at molecular, cellular and circuit levels that induce long-term modifications leading to DRG neuron hyperexcitability, dorsal horn of the spinal cord (DHSC) central sensitization and pain hypersensitivity (*13*). These similarities between MOR- and FLT3-induced pain sensitization prompted us to test whether FLT3 contributes to molecular and functional changes associated with MIT and MIH.

Given the role of DRG neurons in the control of MOR- and FLT3-dependent hyperalgesia (*7, 13*), we looked at the cellular localization of MOR and FLT3 in lumbar DRG neurons of saline-treated control mice and mice receiving repeated morphine (10 mg/kg, subcutaneous injections, twice daily for 4 days) treatment (referred hereafter as chronic morphine). We used the MOR-mCherry mouse line combined with mCherry immunohistochemistry for MOR and *in situ* hybridization for *Flt3* mRNA. We found that, in control mice, 35 ± 1.6% and 25.5 ± 2.3% of DRG neurons were MOR-or *Flt3*-positive (MOR+ or *Flt3+*), respectively (**Fig. 1A, B**). A subset of 13 ± 1.6% of MOR+ neurons were *Flt3*+ and 18.7 ± 3.6% of *Flt3+* neurons expressed MOR (**Fig. 1B**). After chronic morphine exposure, the percentage of MOR+ and *Flt3*+ neurons did not change significantly (41.6 ± 1.6% and 21 ± 1.8%, respectively), whereas the subset of *Flt3*+ neurons expressing MOR was doubled to 34.9 ± 2.3% compared to control conditions (**Fig. 1A, B**). We next investigated the effect of FLT3 genetic inhibition in the development of OIH. Chronic morphine elicited a progressive decrease of the mechanical nociceptive threshold (i.e. indicative of MIH) in both male and female *Flt3*^WT^ littermate mice (**Fig. 1C, S1A)**. By contrast, *Flt3*^*KO*^ littermates failed to develop MIH irrespective of gender (**Fig. 1C, S1A**), which indicates that FLT3 is necessary for the establishment of MIH.

**Fig. 1.**
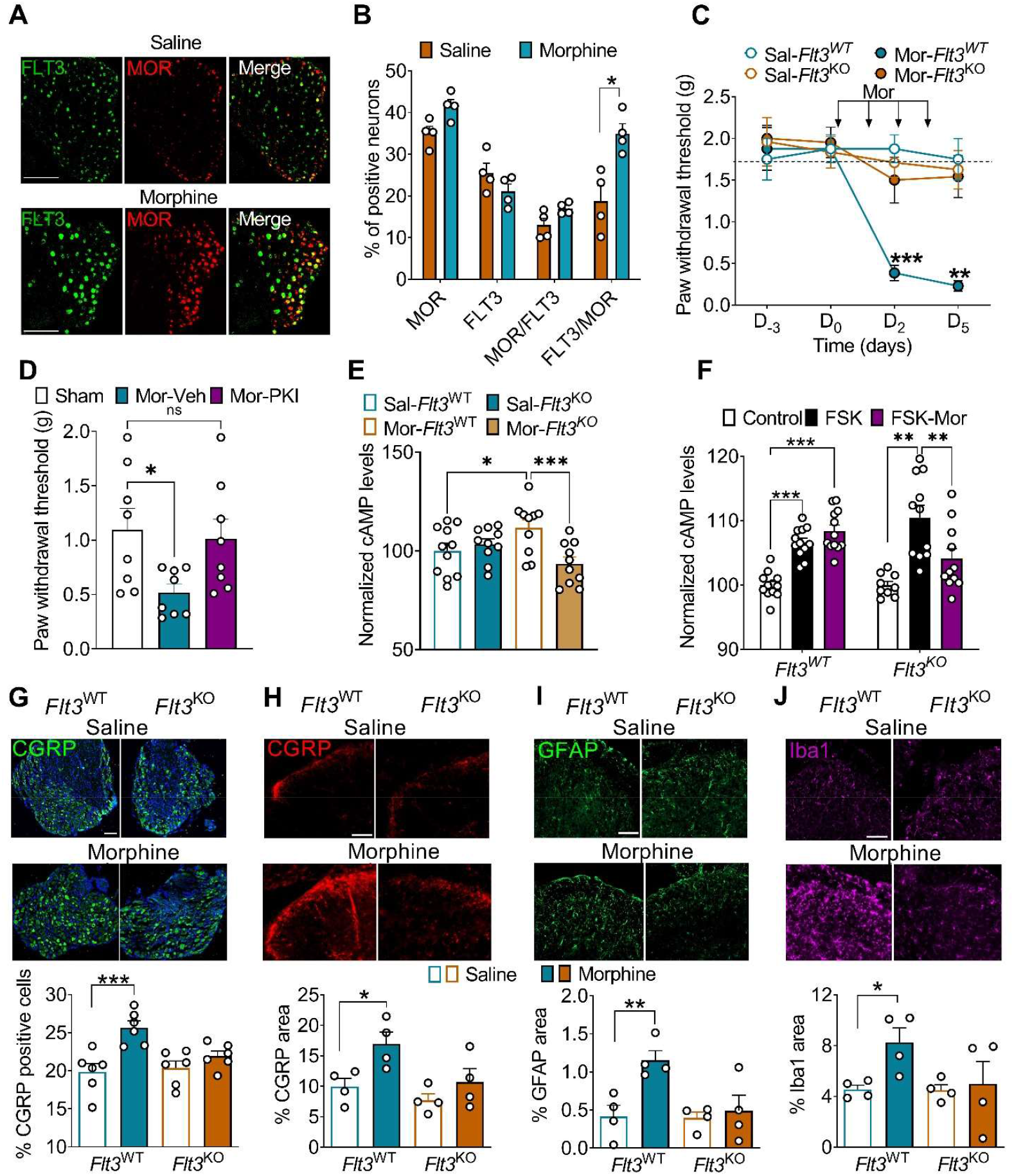
*Flt3* and MOR colocalization in sensory neurons is enhanced by chronic morphine and *Flt3* invalidation blocks MIH and associated signaling and biochemical changes. (**A**) *In situ* hybridization of *Flt3* mRNA in DRG sensory neurons from MOR mCherry mice showing the colocalization of MOR and FLT3 in control mice (top) and mice repeatedly treated with morphine (10 mg/kg i.p., twice a day for 4 days, bottom), respectively. Bars = 100 µm. (**B**) Quantification of the images shown in (A) as number of positive neurons. MOR and FLT3 are the percentage of MOR+ and FLT3+ neurons, respectively, among all DRG neurons. MOR/FLT3 and FLT3/MOR are the percentages of MOR+/FLT3+ co-expressing neurons that are also FLT3+ and MOR+, respectively. (**C**) Mechanical pain sensitivity of *Flt3*^WT^ or *Flt3*^KO^ mice (n = 6) assessed by von Frey after chronic morphine (10mg/kg i.p. twice a day for 4 days,) or chronic saline treatment. **(D)** Mechanical pain sensitivity of *FLT3*^WT^ mice (n = 8) assessed by von Frey test two days after starting chronic morphine with or without PKI (5 µg i.t.). (**E**) cAMP dosage in DRGs from *Flt3*^WT^ or *Flt3*^KO^ mice treated with saline or morphine (n = 10 or 11). (**F**) cAMP responses to forskolin in primary cultures of DRG sensory neurons with morphine from *Flt3*^WT^ (n = 4) or *Flt3*^KO^ (n = 3) mice treated. (**G-J**) **Top**: CGRP-immunoreactivity in DRG sensory neurons and CGRP-, GFAP- and Iba1-immunoreactivities, respectively, in DHSC from *Flt3*^WT^and *Flt3*^KO^ mice (n = 6) after chronic morphine or chronic saline treatment. **Bottom**: quantification of images shown in **G-J**. Results are mean ± SEM of data collected from 4-6 animals. FSK, forskolin; Mor, morphine; Sal, saline. *P < 0.05, **P < 0.01, ***P < 0.001. Statistical analyses included Mann-Whitney test (**B**), two-way repeated ANOVA followed by Bonferroni’s test for multiple comparisons (**C, G-J**) and mixed-effect model (REML) test followed by Šidák’s test for multiple comparisons (**D, E**).

To gain insights into FLT3 mechanisms, we evaluated the effects of FLT3 on morphine-induced overactivation of cyclic adenosine 3′, 5′-monophosphate (cAMP), a canonical and critical second messenger of MOR to generate MIH (*14-16*). Firstly, PKI 14-22, an inhibitor of protein kinase A (*14*), the main effector of the adenylyl cyclase (AC)/cAMP pathway, prevented MIH in mice, confirming the role of cAMP signaling pathway in adaptive changes produced by chronic morphine *in vivo* (**Fig. 1D**). Secondly, cAMP levels were elevated in DRG collected from chronic morphine *Flt3*^WT^, but not from *Flt3*^*KO*^ mice compared to saline *Flt3*^WT^ group (**Fig. 1E)**. Lastly, we performed primary cell culture of adult sensory neurons and evaluated the effects of acute or chronic morphine on MOR-induced cAMP production. As previously reported (*15–17*), acute morphine inhibited forskolin-stimulated cAMP levels in DRG primary cultures from *Flt3*^*WT*^ mice, a process dependent upon the activation of MOR as shown by the inhibitory effects of CTAP, a MOR-selective antagonist, on morphine-induced inhibition of cAMP production (**Fig. S2A**). The inhibitory effect of morphine on forskolin-induced cAMP accumulation was totally absent in primary cultures of DRG neurons chronically exposed to morphine (exposure to 10 µM for 7 days) (**Fig. 1F**). However, the morphine inhibitory effect was maintained when primary cultures were prepared from *Flt3*^*KO*^ mice (**Fig. 1F**). This was unlikely due to a difference in the expression of MOR receptors in the DRG, since no difference was observed between *Flt3*^WT^ and *Flt3*^*KO*^ animals (**Fig. S2B**). These *in vitro* and *in vivo* results indicate that FLT3 is involved in cAMP-dependent pronociceptive changes produced by chronic morphine.

MIH and MIT are adaptive processes that have been proposed to result from complex alterations at the molecular level for MORs, as well as at the cellular and circuit levels, in both the peripheral and central nervous systems (*7*). The increase in CGRP in DRG neurons and glial activation in the DHSC are described as among the main alterations caused by chronic morphine treatment, which results in hyperexcitability of sensory nerve fibers and increased central neurotransmission (*2, 3, 18, 19*). Hence, the role of FLT3 was further investigated by looking at the influence of *Flt3* deletion on these chronic morphine-induced pronociceptive phenotypes. An increase in CGRP immunostaining was observed both in DRG neurons (**Fig. 1G**) and their DHSC projections (**Fig. 1H**) from *Flt3*^WT^ mice after chronic morphine, but was absent in *Flt3*^KO^ mice. Glial changes as evidenced by increase in the expression of GFAP in astrocytes and Iba1 in microglia, were also FLT3-dependent, since they were largely suppressed in *Flt3*^*KO*^ mice (**Fig. 1I, J**). Hence, our data show that FLT3 promotes chronic morphine-induced biochemical changes associated with MIH at both DRG and DHSC levels.

We next asked whether FLT3 also regulates opiate-induced hyperexcitability of sensory neurons typically seen in MIH and MIT. We first used video microscopy intracellular Ca^2+^ ([Ca^2+^]_I_) fluorescence imaging on primary cultures of adult DRG neurons to assess neuronal excitability. In control conditions, DAMGO (10 µM), a MOR-selective agonist which by binding on MOR reduces neuronal excitability by inhibiting voltage-activated Ca^2+^ channels in sensory neurons (*20*), significantly decreased [Ca^2+^]_i_ transients induced by the application of a high K^+^ concentration (HiK^+^; 50 mM) to a similar extent in DRG neurons from *Flt3*^WT^ and *Flt3*^*KO*^ mice (**Fig. 2A, B**). In contrast, after chronic morphine exposure of the DRG neurons (10 µM twice a day for 4 days before [Ca^2+^]_I_ experiments), application of DAMGO increased HiK^+^-induced [Ca^2+^]_I_ transients in *Flt3*^WT^ DRG neurons (**Fig. 2A, B**), confirming MOR excitatory effects upon repeated stimulation (*20*). The DAMGO potentiating effect was absent in neurons from *Flt3*^KO^ mice after chronic morphine treatment (**Fig. 2A, B**), suggesting that *Flt3* deletion prevents the development of morphine-induced DRG neuron hyperexcitability. We also investigated the effects of the FLT3-specific negative allosteric modulator, BDT001, which has been shown to inhibit FL-induced FLT3 phosphorylation and abrogate nerve injury–induced neuropathic pain (*13*). Similarly to *Flt3* genetic deletion, FLT3 pharmacological blockade by BDT001 (1 µM), reversed the HiK^+^-induced [Ca^2+^]_I_ transient potentiation induced by DAMGO in chronic morphine-treated DRG neurons (**Fig. S3A, B**). We recorded single nerve fiber electrical activity using the *ex-vivo* glabrous skin-saphenous-nerve preparation (**Fig. 2C**). Aberrant post-discharge activity was common in sensory fibers from morphine-treated *Flt3*^WT^ mice after skin mechanical stimulation as previously described (*21*), but rare or absent in fibers from morphine-treated *Flt3*^KO^ mice (**Fig. 2D**). Sensory fibers from morphine-treated *Flt3*^WT^ mice continued to respond long after the termination of the mechanical stimulus. Consequently, the mean firing activity of mechano-sensory fibers following mechanical stimulation was significantly increased by 85% in *Flt3*^WT^ chronic morphine mice (**Fig. 2E**). This largely accounted for by an increase in activity in both high threshold mechanonociceptors (Tonic) (+106 ± 21.8%) and low-threshold slowly-adapting (SA) mechanoreceptors (+62 ± 8%). Low-threshold rapidly-adapting (RA) mechanoreceptors showed no obvious aberrant post-discharges following chronic morphine treatment. Remarkably, no post-discharges were observed in mechanosensory nerve filers from chronic morphine *Flt3*^KO^ mice after mechanical stimulation. Here again, BDT001 (5 mg/kg), administered *in vivo* 90 min before each morphine administration, also abolished morphine induced-aberrant post-discharges of both high-threshold mechanonociceptors and low-threshold mechanoreceptors recorded using the skin-saphenous-nerve preparation (**Fig. S3C**). We finally investigated the effects of chronic morphine at DRG central terminal synapses in DHSC, by recording evoked-excitatory postsynaptic currents (eEPSCs) in lamina I and II outer (IIo) neurons of DHSC, in response to electrical stimulation of sensory afferents (**Fig. 2F**). In *Flt3*^WT^ mice, chronic morphine induced a significant increase in the peak amplitude of eEPSCs when compared to chronic saline-treated *Flt3*^WT^ mice (**Fig. 2G, H**). The effect of morphine on eEPSC amplitude was suppressed in *Flt3*^*KO*^ mice. Altogether, these functional data suggest a key role of FLT3 in the development of morphine-induced hyperexcitability along DRG fibers from peripheral to central synapses.

**Fig. 2.**
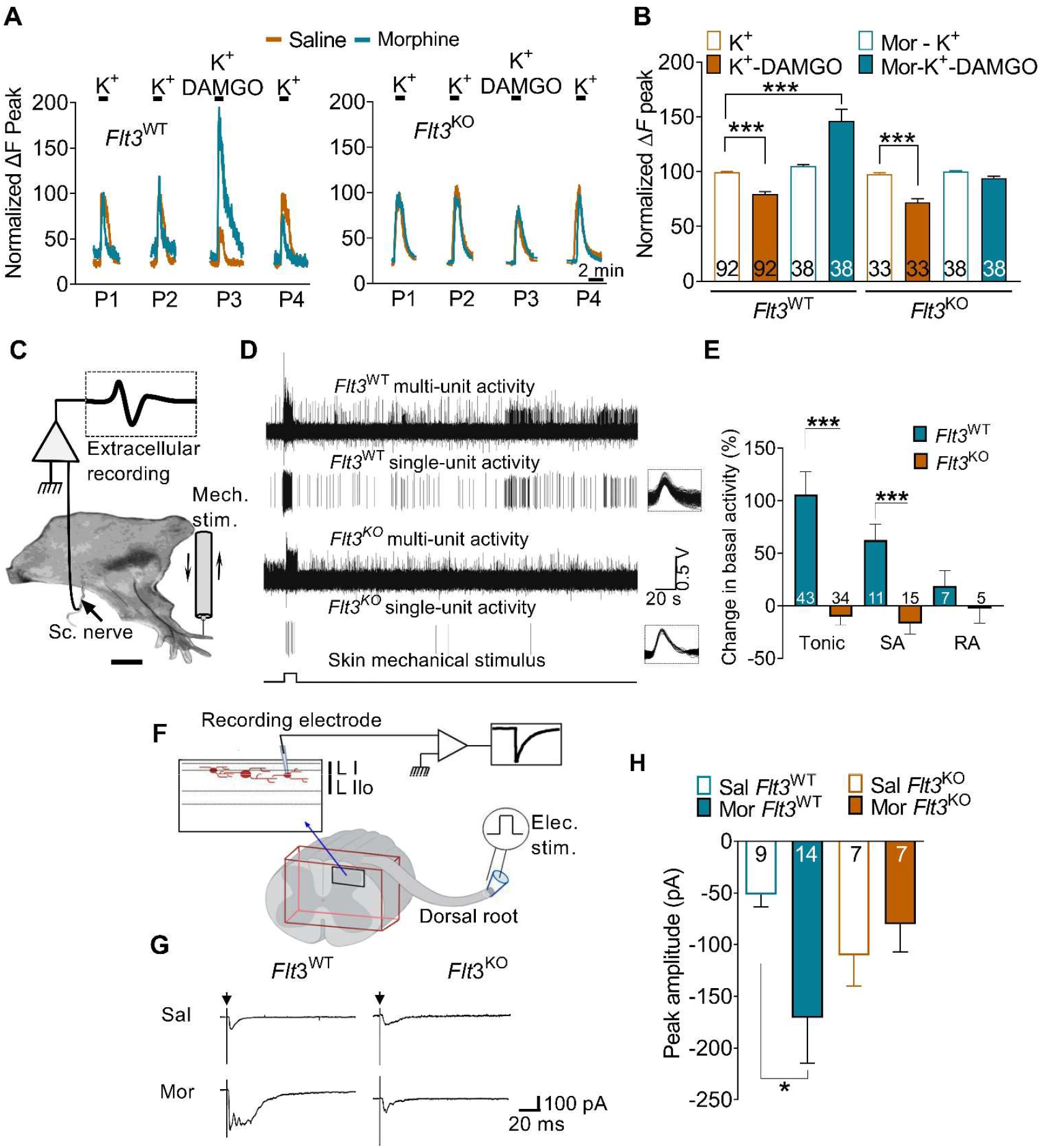
FLT3 controls chronic morphine-induced peripheral neuronal hyperexcitability. **(A)** Traces of [Ca^2+^]_i_ responses to repeated bath applications of high K^+^ (HiK^+^, 50 mM) alone or combined with DAMGO (1 µM), a MOR-selective agonist, in chronic saline or morphine-treated cultured DRG neurons from *Flt3*^WT^ (n = 4) or *Flt3*^*KO*^ (n = 4) mice (morphine at 10 µM for 3-4 DIV, followed by rinsing out). (**B**) Results are expressed as response amplitudes normalized to HiK^+^ alone (ΔF peak). (**C**) Scheme of the skin-nerve preparation set-up, allowing recording of single or multi-unit activities. The mini-chamber allows mechanical stimulation of the selected receptive fields. (**D**) *Ex vivo* extracellular recordings of spontaneous- and mechanically evoked-activity on high threshold C-mechanoreceptors from saphenous-nerve preparations of *Flt3*^WT^ and *Flt3*^KO^ mice before and after a 10 s-mechanical stimulus (*bottom trace*) applied on chronic morphine-treated mice. The extracellular recodings, as well as the single-unit activity recordings derived from the raw action potentials (APs) (*individual traces, rightmost panels*) highlight the presence of post-discharges in *Flt3*^WT^, but not *Flt3*^KO^ mice. (**E**) Normalized firing of cutaneous mechanoreceptors recorded in *Flt3*^WT^ (n = 7) or *Flt3*^KO^ (n = 4) mice exposed to chronic morphine (10 mg/kg ip, twice a days for 4 days. (**F**) Scheme of the electrophysiological set-up to record evoked responses in lamina I-IIo in the DHSC after electrical stimulation of the dorsal root. (**G**) Examples of monosynaptic evoked responses induced by dorsal roots electrical stimulation in spinal slices of *Flt3*^WT^ and *Flt3*^KO^ mice treated with either chronic saline or morphine. (**H**). Peak amplitude of monosynaptic evoked responses of neurons recorded from saline-treated *Flt3*^WT^ (n = 6) or *Flt3*^KO^ (n = 6) mice, and morphine-treated *Flt3*^WT^ (n = 8) or *Flt3*^KO^ (n = 5) mice. Elec. stim, electrical stimulation; Mech. stim, mechanical stimulation; Mor, morphine; Sal, saline; Sc. Nerve, sciatic nerve; Tonic, high threshold mechanonociceptors; RA, low-threshold rapidly-adapting mechanoreceptors; SA, low-threshold slowly-adapting mechanoreceptors.. Bar and symbols in B, E and H represent mean ± SEM of n neurons per group. Statistical analyses included mixed-effects model (REML) followed by the Bonferroni’s test for multiple comparisons (**B, E**) and the Kruskal-Wallis test followed by the Dunnett’s test for multiple comparisons (**H**).

We next investigated whether specific FLT3 impairment, i.e. downregulation of its expression at the DRG neuron level, affects the development of MIH and MIT by using an *Flt3*-targeted shRNA delivered by an intrathecally-injected AAV9 virus (**Fig. 3A-C**). This *Flt3*-shRNA AAV9 virus downregulated FLT3 expression in the DRG (*13*) and blocked FL-induced mechanical pain hypersensitivity without affecting motor function in mice (*13*) and rats (**Fig. S4A, C**). Analgesia tolerance to morphine (**Fig. 3B**), mechanical pain hypersensitivity (**Fig. 3C**) and mechanical allodynia (**Fig. S3B**) were fully prevented in rats receiving a single injection of AAV9 *Flt3*-shRNA, but not the control AAV9 non-targeted shRNA. Finally, FLT3 stimulation by intrathecally injected FLT3 ligand FL (138 nM/10 µL) largely inhibited acute morphine analgesia in intact animals (**Fig. 3D**), demonstrating that FLT3 activation, as does its inhibition, controls MOR function. Altogether, these results support a key role of neuronal FLT3 in the DRG in promoting MIT and MIH.

**Fig. 3.**
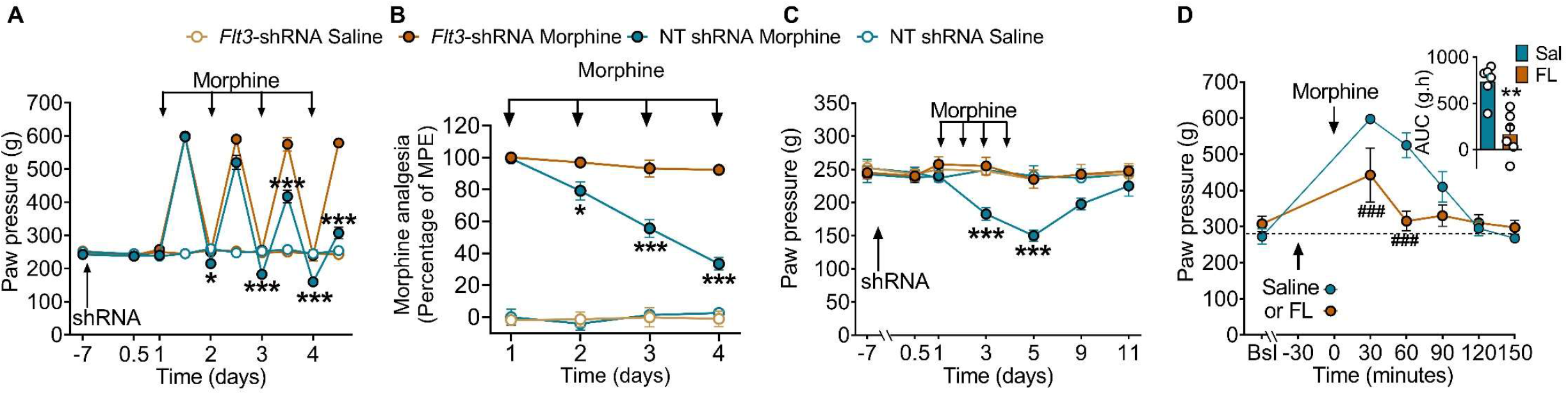
Peripheral FLT3 is required for the onset of MIT and MIH. (**A**) Mechanical pain sensitivity of rats receiving a single intrathecal injection of AAV9 non-target (NT shRNA) shRNA or AAV9 *Flt3*-shRNA virus before and during chronic morphine (2mg/kg twice a day for 4 days, subcutaneously) (n = 5-6 rats/group). (**B**) Representation of anti-nociception as the percentage of Maximal Possible Effect (MPE) after the first daily injection of morphine in AAV9 non-targeted (NT shRNA) shRNA-or AAV9 *Flt3*-shRNA-treated rats. **(C)** Mechanical pain sensitivity of rats receiving a single i.t. injection of AAV9 non-targeted shRNA or AAV9 *Flt3*-shRNA virus before daily morphine treatment. **(D)** Mechanical pain sensitivity of rats receiving a single intrathecal injection of FL (138 nM/20 µL) 30 minutes before an acute morphine injection (2 mg/kg, subcutaneously) (n = 6 rats/group). Inset: Bar and symbols represent mean of area under the curve (AUC). Data are shown as mean ± SEM *P < 0.05, **P < 0.01, ***P < 0.001 vs. NT shRNA; ### P <0.001 vs. Saline. Statistical analyses included mixed-effects (REML) (A) or two-way ANOVA (B-D) followed by the Bonferroni’s test for multiple comparisons and Mann-Whitney test (D, *inset*).

We then asked whether the FLT3 specific inhibitor BDT001 and morphine, when combined, achieved better analgesia than morphine alone. Firstly, BDT001 (5 mg/kg, intraperitoneal) 90 min before subcutaneous morphine (2.5 mg/kg, subcutaneous) treatment significantly potentiated acute morphine analgesia in rats, as shown by the leftward shift in the morphine dose-response curve, which represents a ∼30% morphine-sparing effect (**Fig. 4A, B)**. Similar results were obtained in mice (**Fig. S5A**). We also showed that intraperitoneal injections of BDT001 totally prevented MIH and MIT in chronic morphine-treated rats (**Fig. S5B, C**). In addition, BDT001 alone did not produce respiratory depression nor constipation and did not influence their intensity nor duration when produced by chronic morphine (**Fig. 4C, D**). Also, BDT001 did not change the withdrawal signs elicited by the opiate antagonist naloxone in chronic morphine (*22*) (**Fig. S5D**). In the conditioned place preference paradigm, BDT001 did not induce motivational effects, nor did it reveal these effects in animals receiving a non-rewarding low dose of morphine (**Fig. S5E**). Hence, the augmentation of morphine analgesia by BDT001 was not accompanied by an aggravation of morphine adverse effects.

**Fig. 4.**
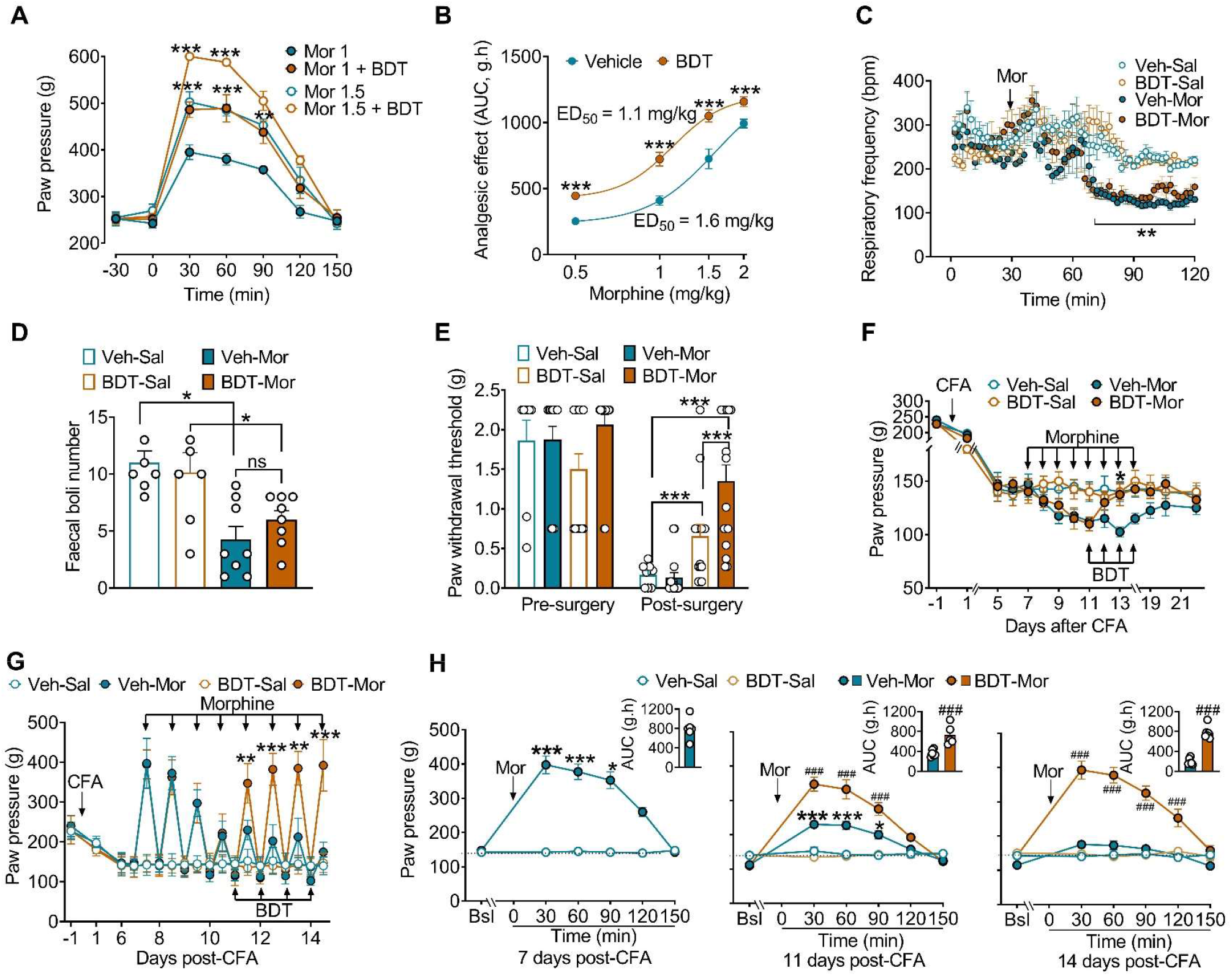
Pharmacological inhibition of FLT3 by BDT001 improves morphine analgesia in normal rats or in acute post-surgical and chronic inflammatory pain without affecting morphine side-effects. (**A**) Dose-response curve of morphine (Mor) analgesia with or without pre-treatment with intraperitoneal (i.p.) BDT001 (BDT, 5 mg/kg) injected 90 minutes before subcutaneous (s.c.) morphine administration. (**B**) Dose-response curve represented by the AUC quantification of morphine analgesia as shown in (A) (n = 5-6 rats/group). (**C**) Respiratory frequency in WT mice after s.c. saline or morphine injection (10 mg/kg pretreated 90 min before with i.p. vehicle or BDT001 (5 mg/kg) (n= 8 mice/group). (**D**) Constipation, measured as the weight of foecal boil in mice receiving i.p. vehicle or BDT001 (5 mg/kg) and then s.c. saline or morphine (10 mg/kg) injection (n= 8 mice/group), 90 min later. (**E**) Effect of i.p. BDT001 (5 mg/kg) alone or in combination with s.c. morphine (1.5 mg/kg) on post-surgical pain (n= 8-14 mice/group). Paw withdrawal thresholds were measured by von Frey before (pre-surgery) of 24 h after surgery (post-surgery). Saline or morphine were administered 24 h after surgery, 30 min before pain assessment. (**F**) Curative effect of i.p. BDT001 (5 mg/kg) on MIH in CFA-treated rats (n= 6 rats/group). BDT001 treatment was initiated on the 5^th^ day of morphine treatment, which was continued for 3 more days. Mechanical pain sensitivity was assessed as in A before morphine treatment. (**G**) Morphine antinociception (percentage of MPE) at the first daily injection of morphine alone or in combination with vehicle or BDT001. (**H**) Time-course and AUC analysis (inset) of the morphine analgesic effect combined with vehicle or BDT001 administration at 7, 11 and 14 days after CFA. Inset: Bar and symbols represent mean of AUC ± SEM of N rats per group. *P < 0.05, **P < 0.01, ***P < 0.001 vs. morphine alone (A-C, F, G); *P < 0.05, ***P < 0.001 for the indicated comparisons (D, E); *P < 0.05, ***P < 0.001 vs. Vehicle (Veh)-Saline (Sal) and ### P < 0.001 vs. morphine alone. Statistical analyses included two-way ANOVA (A-C, E-H) or mixed-effects (REML) (D) followed by the Bonferroni’s test for multiple comparisons and Mann-Whitney test (H, *insets*).

We next tested BDT001, alone or in combination with morphine, in two pathophysiological pain animal models. Using the incisional post-surgical pain model in mice (*23, 24*), we found that at 24 h post-surgery the nociceptive threshold was strongly decreased to a similar extent in control and low morphine dose-treated animals, indicating that the dose of morphine used (1.5 mg/kg) was ineffective (**Fig. 4E**). BDT001 alone, administered during the surgical procedure, produced a partial but significant analgesic effect. Interestingly, morphine administration (1.5 mg/kg) to BDT001-treated mice displayed a marked increase in the nociceptive threshold (**Fig. 4E**), indicating that a synergistic drug combination could allow lower doses of morphine to be efficient. Moreover, the BDT001-morphine drug combination prevented MIH and MIT in the Complete Freund’s Adjuvant (CFA) model of persistent inflammatory pain (*25*). As classically described, CFA injection in the hindpaw induced a decrease in mechanical nociceptive threshold (**Fig. 4F, G and S6A, C)**. Animals further receiving chronic morphine show a progressive loss of morphine analgesia, i.e. tolerance (**Fig. 4H, S6B**) and an exaggeration of CFA-induced mechanical pain hypersensitivity (MIH) (**Fig. 4F, S6C**). Concomitant administration of BDT001 with morphine enhanced morphine analgesia and prevented morphine tolerance (**Fig. S6A, B**). The combination of morphine with BDT001 had modest effects on CFA-induced inflammation as evaluated by the diameter of the injected hindpaw (**Fig. S6D**), suggesting that the restoration of morphine analgesia was unlikely due to an action of BDT001 on peripheral inflammation. Additional experiments were conducted to evaluate the curative effects of BDT001 on MIT and MIH (**Fig. 4F-H**). Rats received morphine alone, twice daily for 4 days (7 to 10 days post-CFA), until MIH and MIT had been established, and morphine treatment was then combined with BDT001 in some animals from 11 to 14 days post-CFA. The baseline nociceptive threshold, measured each day before the first morphine treatment, was progressively normalized in animals treated with BDT001 (**Fig. 4F**), indicative of a reversion of MIH. The first injection of morphine (7 days post-CFA) produced a robust analgesic effect that decreased over the time with the repetition of the injections, i.e. once MIT was installed (**Fig. 4G, H)**. Importantly, as soon as the first BDT001 injection was performed (11 days post-CFA), morphine analgesia was increased and entirely restored after 4 days of BDT001 treatment (15 days post-CFA) (**Fig. 4G, H**). Altogether, these results indicate that BDT001 therapy, in association with morphine, improves acute and chronic morphine analgesia in pain models by preventing or reversing MIT and MIH. This therapy has, therefore, the potential of limiting morphine dose escalation, and thereby life-threatening adverse effects in humans.

Our study provides new evidence that FLT3 functioning in primary sensory neurons is critically involved in both opioid-induced analgesia and chronic opioid-induced MIT and MIH. Along with the alteration of MOR-dependent effects by FLT3 activation or its genetic and pharmacological inhibition, our results show that FLT3 regulates MOR signaling through the canonical cAMP-PKA pathway that produces neuronal hyperexcitability. These observations suggest a functional interaction between MOR and FLT3 receptors in sensory neurons, driving the initiation of adverse counter-adaptations to opioid analgesia, and allowing the onset of MIT and MIH. How MOR and FLT3 interact at the molecular level remains to be determined. Additionally, this study provides a scientific basis to develop therapeutic strategies for disrupting FLT3 signaling to maintain adequate morphine pain relief under chronic morphine exposure. Indeed, BDT001, a new specific negative allosteric modulator of FLT3 (*13*), improves morphine analgesia and prevents morphine tolerance in persistent inflammatory pain. In addition, BDT001 alone or in combination with morphine does not elicit or worsen the morphine adverse effects. These side effects are positively correlated with opioid dose escalation, leading to an increase of morbidity and mortality, underlying the present Opioid Crisis (*5, 26*). Thus, FLT3 inhibitors, which avoid dose escalation without decreasing opioid analgesia, show some promise as agents to improve chronic opioid analgesic therapy. In conclusion, our fundamental findings could have important clinical implications, raising the exciting possibility that effective treatments might be available soon for the untold millions of patients suffering from chronic pain.

## Acknowledgments

We thank Dr Lemischka (The Black Family Stem Cell Institute, Icahn School of Medicine at Mount Sinai, New York) for providing the *Flt3* knock-out mice. We thank Dr Dominique Massotte (UPR3212 Institut des Neurosciences Cellulaires et Intégratives (INCI)) for providing the MOR-mCherry mice. We also thank the different technical platforms of Institut des Neurosciences de Montpellier (INM), especially the animal care facility and the functional exploration platform RAM Neuro supervised by Denis Greuet, and RIO imaging platform supervised by Chamroeur Sar.

## Funding

This work was supported by INM, INSERM, CNRS and Aix-Marseille and Montpellier University and grants from:

National Research Agency (grant ANR-19-CE16-0008),

Fondation pour la Recherche Médicale (DEQ20130326482 to PD), CBS2 doctoral school,

French ministry of higher education, research and innovation

Biodol Therapeutics, France.

## Author contributions

Conceptualization: JV, CR

Methodology: MA, RD, PD, NF, IM, JR, JV, CR

Investigation: CR, AT, MT, MA, JR, LD, AJ, JPL, FD, SM, JB, CS, CS, SV, IM

Funding acquisition: RD, PD, NF, DR, PS, JV, CR

Project administration: JV, CR

Supervision: JV, CR

Writing – original draft: MA, RD, PD, AT, MT, JR, PC, CR, PS, JV

Writing – review & editing:

