## Supplementary Materials for "FLT3 signaling inhibition preserves opioid analgesia while abrogating tolerance and hyperalgesia"

Adresses: [1]Université de Montpellier, Montpellier, France ; [2]Inserm U-1298, Institut des Neurosciences de Montpellier, Montpellier, France ; [3]Laboratoire de Neurosciences Cognitives, CNRS AMU, Marseille, France ; [4]NeuroDol INSERM 1107, Clermont-Ferrand, France ; [5]Biodol Therapeutics, Cap Alpha, Clapiers 34830, France, [6]Laboratoire d'Innovation Thérapeutique, UMR7200 CNRS/Université de Strasbourg and LabEx MEDALIS, Faculté de Pharmacie, 74 route du Rhin, 67412 Illkirch, France.

$These authors contributed equally: Antoine Jouvenel, Adrien Tassou, Maxime Thouaye.

Materials and Methods

Figs. S1 to S6

References (*27-31*)

### Materials and Methods

#### Animals

Experiments were performed in male Sprague-Dawley rats (150-170g, Janvier, France), C57BL/6 naive mice (25-30g, Janvier, France), or C57BL/6 mice carrying a homozygous deletion of Flt3 (*Flt3*$^{KO}$ mice) and their littermates (WT) weighing 25-30 g. All the procedures were approved by the French Ministry of Research (authorization #1006). Animals were maintained in a climate-controlled room on a 12 h light/dark cycle and allowed access to food and water ad libitum. For rats and mice, we used n = 6 or 8 per group for each experiment respectively, otherwise the number is noted in the legends. Male and female mice were first considered separately in behavioral procedures. Both sexes showed mechanical hypersensitivity of same intensity after repeated morphine injection and were similarly affected by Flt3 deletion (ANOVA followed by Bonferroni's test, n = 8 for both sexes and genotypes for each experiment, Fig. S1). Thereafter, experiments were performed only on male mice.

#### Drug delivery

*Opioids*. Morphine sulfate was purchased from Francopia, dissolved in 0.9% sodium chloride and administered subcutaneously. For the chronic morphine administration protocol, morphine subcutaneous injections were performed twice a day with 8h of interval during 4 days. Rats and mice received 2 mg/kg and 10 mg/kg per injection, respectively. For acute morphine-FL treatment protocol in rats (**Fig. 3D**), one subcutaneous morphine injection (2 mg/kg) was performed 30 minutes after FL (138nM/50µl) intrathecal injection.

*FL*. Human recombinant FL (FLT3 ligand) was obtained in the E. coli Rosetta (DE3) strain (Novagen) in our laboratory using the pET15b-rhFL plasmid and was dissolved in saline solution. For recombinant FL intrathecal injection, a 30 G needle attached to a Hamilton syringe was inserted between L4 and L5 vertebrae in lightly restrained, isoflurane-anaesthesized rats. The reflexive tail flick was used to confirm the punction. A total volume of 10-20 µl was injected.

*Pharmacological inhibitors*. Different inhibitors were used for the *in vivo* experiments in rats. Intrathecal injections were performed as previously described for FL, 1h before intrathecal injection of FL, the PKA inhibitor PKI 14-22 (#2546 – Tocris) was administered at a concentration of 5µg/20µl.

*BDT001*. BDT001 was synthesized by D. Rognan at Strasbourg (Laboratoire d'Innovation Thérapeutique, UMR7200, CNRS – Université de Strasbourg, Illkirch 67400 France). BDT001 was dissolved in 7% Tween 20 in a saline solution with 6 sonication runs of 45 seconds. For chronic morphine protocol with BDT001 treatment, intraperitoneal injection of BDT001 was performed 2h before end of the evening morphine injection. BDT001 was injected at 5mg/kg in mice and at 2mg/kg in rats.

### Pain model

*Inflammation*: The model of complete Freund adjuvant (CFA)-induced pain has been used for assessing chronic inflammatory pain (*25*). Under isoflurane anesthesia, an intraplantar injection (50µL) of a solution of 1mg of mycobacterium tuberculosis (Sigma-Aldrich) per ml was performed in the right hindpaw of wild-type Sprague-Dawley rats. Once CFA-induced hypersensitivity is established, chronic morphine administration was performed as described above. For preventive experiments, morphine and BDT001 administration started on the same day for 4 days. For the curative experiments, chronic morphine was first administered and once analgesic tolerance and morphine-induced hyperalgesia was established, BDT001 was started with the following morphine injections.

*Paw incision models*: Mice C57BL/6J were anesthetized under isoflurane (3% vol/vol). For the single incision (SI) model (*23*), a 0.7cm incision was applied with a number 20 blade on the skin and fascia of the left hindpaw plantar surface. Subcutaneous plantaris muscle was then isolated, exposed and longitudinally incised. After hemostasis, the wound was sutured with two 6.0 absorbable sutures and the mice were finally placed in recovery cages. BDT001 or vehicle was administered 90 minutes before the incision. On D1, a single saline or morphine injection (1.5 mg/kg; i.p.) was administered and the nociceptive threshold was evaluated 30 minutes later.

### Behavioral analysis

Before testing, rats and mice were acclimatized for 60 min in the temperature and light-controlled testing room within a plastic cylinder or on wire mesh. Experimenters were blinded to the genotype or the drug administered Since acute active morphine injection induce motor hyperactivity in mice but not in rats, we chosed rats to study morphine analgesia in details to obtain a better temporal assessment of the analgesia by repeated (every 30min) measurements of the morphine effect.

*Tactile sensitivity*. Tactile withdrawal threshold was determined in response to probing of the hindpaw with eight calibrated von Frey filaments (Stoeling, Wood Dale, IL, USA) in logarithmically spaced increments ranging from 0.04 to 8 g (4–150 mN). Filaments were applied perpendicularly to the plantar surface of the paw. The 50% paw withdrawal threshold was determined in grams by the Dixon nonparametric test (*27*). The protocol was repeated until three changes in behaviour occurred. This method was used in both rats and mice with adapted filaments.

*Paw pressure test*: To assess mechanical sensitivity in rats, a variant method of the Randall-Selitto method was performed as previously described (*28*). Briefly, increasing pressure was applied to the hind paw until the rat squeaks. The Basile analgesimeter (Apelex, Massy, France; stylus tip diameter, 1 mm) was used. A 600-g cutoff value was determined to prevent tissue damage. After their arrival to the laboratory, rats were kept in their colony room for 4 days. To avoid any experimental stress, the same experimenter in quiet conditions realized experimentations. Animals were habituated to experimenters, experimental room and devices for at least 10 days before starting the experiments. Before starting the experiments, basal nociceptive threshold was measured during 2 days in a row and on the experimental day to verify nociceptive threshold stability. To assess morphine analgesia, measurements were performed 30 minutes after morphine injection and in some experiments, the time course of morphine analgesia was determined by measuring nociceptive threshold every 30 minutes for 2h30 until return to a basal nociceptive threshold. To assess morphine induced pain hypersensitivity (MIH), basal nociceptive threshold was assessed once daily before twice daily morphine administration or after FL-morphine administration.

*Morphine and BDT001 motivational effects*. To evaluate morphine or BDT001 motivational effects, a 5 days CPP protocol was set. The apparatus (Bioseb, France) consists in 2 chambers (size 20 cm x 18cm x 25 cm) distinguished by the texture of the floor and by the wall patterns connected to each other by a central chamber (size 20 cm x 7 cm x 25 cm). During the first day, all the mice were able to move freely during 15 minutes in the whole apparatus. Their moves were recorded with a camera connected to Ethovision software. Mice with a spontaneous preference up to 75% were removed from the experiment. The next 3 days are the conditioning days. The morning, mice were restricted for 20 minutes to one chamber after morphine (1.5 or 5 mg/kg) or BDT001 (5 mg/kg) intraperitoneal injection. The afternoon, mice were again restricted for 20 minutes to the opposite chamber after vehicle intraperitoneal injection. Injections were switched off the next day in order to avoid time dependent conditioning. To evaluate the effects of BDT001 on morphine-induced motivational effects BDT001 or vehicle was administered in combination with morphine 2 hours before morphine or saline conditioning. The 5th day is the test day. Mice were placed in the center chamber with free access to all chambers and the time spent in each chamber was recorded for 15 minutes. The preference scores were expressed as the difference between the time spent in the morphine- and/or BDT001-paired chamber and the vehicle-paired chamber.

*Morphine side effects*: Respiratory depression measurement using whole body plethysmography Ventilatory parameters were recorded as previously described (*27*) in conscious C57BL/6N mice by whole-body barometric plethysmography (Emka Technologies, Paris, France). Mice were acclimatized with the plethysmograph chamber for 30 minutes until a stable baseline was obtained. Then, the animal was gently removed from the chamber for subcutaneous injection of the tested drug at T0 and replaced in the chamber for the remaining measurements. Respiratory frequency (f) was recorded for 100 minutes and used as the index of respiratory depression. Accumulated faecal boli quantification was performed as previously described (*29*). Mice were subcutaneously injected with vehicle or BDT001 (5 mg/kg) and 1.5h later they received saline or morphine (10 mg/kg) and individually placed into small Plexiglas boxes lined with filter paper. Faecal boli were collected and weighed every hour for 5h after the drug injections.

*Physical dependence studies*. After induction of chronic tolerance, withdrawal was precipitated by 1 mg/kg naloxone hydrochloride injection performed 2h after the last injection of morphine. Mice were individually placed in small Plexiglas boxes and observed and scored for 20 min for manifestation of different withdrawal signs, including jumping, grooming, rearing, forepaw shaking, genital licking and wet-dog shakes. A global withdrawal score, excluding weight loss, was calculated as previously described (*22*). The sum of all weighted signs produced a global withdrawal score for each mouse.

*Statistical analysis*. All experiments were randomized. Data are expressed as the mean ± SEM. All sample sizes were chosen based on our previous studies except for animal studies for which sample size has been estimated via a power analysis using the G-power software. The power of all target values was 80% with an alpha level of 0.05 to detect a difference of 50%. Statistical significance was determined by analysis of variance (ANOVA one-way or two-way for repeated measures, over time), followed by Bonferonni's post-hoc test for multiple comparisons. To quantitatively evaluate morphine analgesic tolerance (MIT), the % maximal possible effect (%MPE) was evaluated by normalization of nociceptive threshold values during analgesia on Randall-Selitto cut-off value, as following: [ (nociceptive threshold value 30' after morphine injection) – (basal nociceptive threshold) ] ÷ [ (cut-off value) – (basal nociceptive threshold) ]. To quantitatively evaluate morphine analgesia, the area under the curve (AUC) was determined as following. The surface areas were calculated by summing the nociceptive threshold values measured every day after the planned day of experimentation, as follows: Surface area = Σ (nociceptive threshold values at H + n) − basal value × n, where n is the number of intervals. This total value is proportional to the surface area because the intervals between successive tests were similar (30 min). The results were expressed as a mean percentage (±SEM) of the reference AUC defined as the AUC calculated in morphine group alone (100% = AUC of morphine group). In this representation with unpaired data and without repeated measures, Mann-Witney tests were performed to evaluate results statistical significance. To determine EC50 of morphine in presence of Vehicle or BDT001, a sigmoidal dose-response curve was generated with GraphPad Prism software (GraphPad Software, Inc., San Diego, USA) based on AUC raw data.

**Virus transfection**.

The following oligonucleotides were used to build the AAVGFP-shRNA plasmids: shFLT3 (GATCGGTGTCGAGCAGTACTCTAAATCAAGAGTTTAGAGTACTGCTCGACACCTTTTT(top)); AGCTAAAAAGGTGTCGAGCAGTACTCTAAACTCTTGATTTAGAGTACTGCTCGACACC(bottom);Sigma-Aldrich (TRC00000378670); non-targeted sh (GATCCAACAAGATGAAGAGCACCAATCAAGAGTTGGTGCTCTTCATCTTGTTGTTTTT (top)); AGCTAAAAACAACAAGATGAAGAGCACCAACTCTTGATTGGTGCTCTTCATCTTGTTG (bottom).The oligonucleotides were annealed and cloned BamHI–HindIII in a vector containing the U6 promoter sequence and the hGH polyadenylation sequence upstream and downstream, respectively, of the BamHI site. The U6-shRNA-hGH sequences were then cut using PmlI and HpaI and cloned in the pAAV-CMV-turbo GFP vector (Cell Biolabs Inc.) linearized with PmlI. All the constructs were checked by DNA sequencing. The viruses were produced by the Viral Vector Production Unit of Institut de la Vision, Paris, France. For in vivo experiments, 10 µl of AAV-turbo GFP-sh Flt3 or AAV-turbo GFP- sh control solution were intrathecally injected in lightly restrained, unanaesthetized rats, as described above. The titer of the virus solution was 5.8 × 1012 genome copies (gc)/ml.

**Adult sensory neuron culture.**

Neuron cultures were established from lumbar (L4–L6) dorsal root ganglia (*13*). Briefly, ganglia were successively treated by two incubations with collagenase A (1 mg/ml, Roche Diagnostic, France) for 45 min (37°C) and trypsin-EDTA (0.25%, Sigma, St Quentin Fallavier, France) for 30 min. They were mechanically dissociated through the tip of a fire-polished Pasteur pipette in neurobasal culture medium (Life Technologies, Cergy-Pontoise, France) supplemented with 10% fetal bovine serum and DNase (50 U/ml, Sigma). Isolated cells were collected by centrifugation and suspended in neurobasal culture medium supplemented with 2% B27 (Life Technologies), 2mM glutamine, penicillin/streptomycin (20 U/ml, 0.2 mg/ml) plated at a density of 2500 neurons per well in 96-well plates and were incubated in a humidified 95% air-5% CO2 atmosphere at 37°C.

**cAMP activation measurement by *cAMP time-resolved FRET assays*.**

Measurement of cAMP accumulation was performed using the cAMP-Gi immunoassay kits (Cisbio Bioassays). The cAMP assay uses a cryptate-conjugated anti-cAMP monoclonal antibody and d2-labeled cAMP. Cultured adult sensory neuron cells were tested at 8 DIV after seeding in a 96-well plate. Prior to lysis and the addition of the cAMP-cryptate-antibody and cAMP-d2, cells were treated with the indicated test compounds in absence or in presence of Forskolin (FSK) at a concentration of 0.1 µM (to stimulate adenylate cyclase). Briefly, for acute treatment, the cells at 8DIV, were pretreated with the indicated concentrations of CTAP (#C6352 - Sigma-Aldrich) for 20 min at 37°C and stimulated 15 min at 37°C with 10µM of morphine (Francopia) in Stimulation Buffer containing 0.5mM of IBMX (#I7018 – Sigma-Aldrich), prior the addition of 0.1µM of forskolin (#F6886 - Sigma-Aldrich) during 15min at 37°C. The reaction was stopped by lysis buffer containing HTRF® assay reagents: the Europium Cryptate-labeled anti-cAMP antibody and the d2-labeled cAMP. The plate was then incubated for 1 h at room temperature before reading the fluorescence emission at 620 and 665 nm using a PHERAstar FS plate reader (BMG Labtech). In these conditions, acute morphine reduced forskolin-stimulated cAMP levels in DRG primary cultures from $Flt3^{WT}$ mice and CTAP, a MOR-selective antagonist, blocked the inhibitory effects of morphine-induced inhibition of cAMP production by 95.45% ± 2.26 ($n = 11$; $p < 0.05$, one-way ANOVA followed by Bonferroni's test for multiple comparisons), indicating a MOR-dependent mechanism of morphine inhibition.

For chronic treatment, the cells were treated every day with 10 µM of morphine, 24h after seeding in a 96-well plate and during seven days. At 8DIV, the same treatment protocol as that of the acute treatment was used to determine of the cAMP accumulation in DRG neurons.

For the measurement of cAMP accumulation in DRG, mice were treated with the saline buffer or morphine at 10mg/kg in I.P twice a day during 3 days. The last day, Forskolin (FSK) was administrated at 1.5µg intrathecal to stimulate adenylate cyclase. After 30 minutes, DRG (L1-L6) were collected, lysed and homogenized in the MagNA Lyser Instrument (Roche; twice for the speed of 6000 for 30 seconds) with ceramic spheres (Lysing Matrix D, MP Biomedicals, #116913100) in lysis & detection buffer 8 (64CL8FDD - Cisbio Bioassays) supplemented with complete protease inhibitor cocktail (Sigma-Aldrich #P8340) and 0.5mM of IBMX (#I7018 – Sigma-Aldrich). Then, centrifugation (10 000 rpm for 15min at 4°C) was performed. Supernatants were collected and cAMP accumulation was measured as described above.

**Real-Time qRT-PCR**

*Flt3*$^{WT}$ and *Flt3*$^{KO}$ mice lumbar (L4–L6) dorsal root ganglia were dissected. RNA was extracted using the RNAqueous-4PCR Kit (Ambion) according to the manufacturer's protocol. The Real-Time qRT-PCR experiment has been described previously (*13*). Briefly 1 µg of total RNA was reverse-transcribed with 100 U of Superscript II reverse transcriptase (Invitrogen) and real-time PCR was carried using SYBR Green I dye detection on the Light Cycler system (Roche Molecular Biochemicals). The relative amounts of specifically amplified cDNAs were normalized with the geometric mean of RNA polymerase II polypeptide J (*Polr2j*) and DEAD box polypeptide 48 (*Ddx48*) as stable control genes using the delta-CT method. Sequences of the primer pairs used are as follows:

*Polr2j*: F-ACCACACTCTGGGGAACATC, R-CTCGCTGATGAGGTCTGTGA; NM_011293 ; *Ddx48*: F-GGAGTTAGCGGTGCAGATTC, R-AGCATCTTGATAGCCCGTGT; NM_138669 ; *Flt3*: F-ATCCCCAGAAGACCTCCAGT, R- CTGGGTCTCTGTCACGTTCA; NM_010229.2 ; *Oprm1*: F-ACAGGCAGGGGTCCATAGAT, R- AATACAGCCACGACCACCAG; NM_001304948.1

**Calcium imaging on cultured DRG neurons**

For calcium imaging video microscopy [Ca2+]i fluorescence imaging, DRG neurons were loaded with fluorescent dye 2.5 µM Fura-2 AM (Invitrogen, Carlsbad, CA) for 30 min at 37 °C in standard external solution contained: 145 mM NaCl, 5 mM KCl, 2 mM CaCl2, 2 mM MgCl2, 10 mM HEPES, 10 mM glucose (pH adjusted to 7.4 with NaOH and osmolarity between 300 and 310 mOsm). The coverslips were placed on a stage of Zeiss Axiovert 200 inverted microscope (Zeiss, München). Observations were made at room temperature (20-23°C) with a 20X UApo/340 objective. Fluorescence intensity at 505 nm with excitation at 340nm and 380 nm were captured as digital images (sampling rates of 0.1-2 s). Regions of interest were identified within the soma from which quantitative measurements were made by re-analysis of stored image sequences using MetaFluor Ratio Imaging software. [Ca2+]i was determined by ratiometric method of Fura-2 fluorescence from calibration of series of buffered Ca2+ standards. Neurons were distinguished from non-neuronal cells by applying 50 mM KCl (HiK+), which induced a rapid increase of [Ca2+]i only in neurons. Only neurons with resting 340/380 fluorescence intensity ratio of 0.25-0.75 were included. We selected primary sensory neurons with small diameters which are known to be nociceptors and to contain MOR-expressing nociceptor subpopulation. Pulses of (50 mM) were applied at around 2 min-intervals and DAMGO (10 µM) was added after the third pulse. After DAMGO application, another HiK+ stimuli was done to demonstrate recovery of the drug. All drugs and solutions were applied with a gravity-driven perfusion system. For chronic treatment, the cells were treated twice a day with either morphine (10µM) or morphine and BDT001 (1 µM) during at least 72 hours. For data analysis, amplitudes of [Ca2+]i increases, ΔF/Fmax, caused by stimulation of neurons with HiK+ were measured by subtracting the 'baseline' F/Fmax (mean for 30 s prior to HiK+ perfusion) from the peak F/Fmax achieved on exposure to HiK+. In the absence of any treatment, the distribution of these ratios was well fitted by a normal distribution. The DAMGO-induced [Ca2+]I transient were expressed as a percentage of the average peak produced by the two previous HiK+ stimuli. To avoid variations, control and drug treatment values were collected from the same cell. Data were collected from at least three different preparations.

**Immunohistochemistry**

Mice were transcardially perfused with phosphate-buffered saline (PBS). DRG (L4-L6), spinal cord (lumbar segment) were collected and post-fixed in 4% paraformaldehyde between 10 min and 2 h depending on the antibody and tissue before being cryoprotected in 30% sucrose in PBS. Tissues were then frozen in O.C.T (Sakura Finetek). Sections (12 to 14µm) were prepared using the Cryostat Leica CM2800E. For immunostaining, frozen sections were blocked and permeabilized in Ca2+/Mg2+-free PBS containing 10% donkey serum 0.1% Triton x-100 during 30 minutes. The sections were then incubated with primary antibodies, at 4°C overnight in PBS free containing 1% donkey serum and 0.1% Triton x-100. After wash in PBS free (3 times for 10 min minimum each), sections were incubated with appropriate secondary antibody conjugated to AlexaFluor and Hoechst (Sigma 1 µg/ml), in the same buffer as the primary antibodies, at room temperature for 1 hour and then washing (3 times for 15 min each), before mounting with mowiol. Images were acquired under a Zeiss slide scanner Axio Scan Z.1 for quantification and under Zeiss LSM 700 Confocal microscope for illustration with ZEN software (Carl Zeiss Microscopy). The following primary antibodies were employed: anti-CGRP

Abcam ab36001 (goat 1:1000), anti-GFAP Abcam ab4674 (chicken 1:1000), anti-Iba1 Wako (rabbit 1:500). The following secondary antibodies were used anti-rabbit AlexaFluor 555/488/647 (donkey 1:1000), anti-goat AlexaFluor 488 (donkey 1:1000) from Life Technologies.

The quantification of CGRP immunostaining in DRG neurons was realized by counting the number of cells expressing CGRP. Ratio with total neuron number was done in different conditions, and results were expressed in mean of obtained ratios (± SEM) and statistical significance was determined by variance analysis (ANOVA one-way) followed by Bonferroni post-hoc test. The number of neurons expressing the various molecular markers of sensory neurons (subtypes) was determined by counting cells with NeuN immunostaining. A minimum of six sections from lumbar DRG were counted from at least 6 animals for each genotype. For Iba1, GFAP and CGRP the quantification was assessed by determining staining surface among spinal cord dorsal horns with ImageJ software, after setting a density threshold above background. Results were shown in mean (± SEM) and statistical significance was determined by variance analysis (ANOVA one-way) followed by Bonferroni post-hoc test. A minimum of 5 sections of dorsal horn were quantified from at least 4 animals per condition.

#### ***In situ* hybridization**

Sense and antisense digoxigenin (DIG)-labelled RNA probes were generated from a mouse Flt3 cDNA clone (IRAMp995N1310Q, GenomeCUBE Source Bioscience) in a 20 µl reaction containing 1 µg of linearized plasmid (digested with NheI ) using the DIG RNA labelling mix (Roche Diagnostics) and Sp6 RNA polymerases (Promega) following the manufacturer's instructions. DIG-labelled RNA probes were purified on MicroSpin G50 columns (GE Healthcare). Saline or morphine-treated WT and *Flt3*$^{KO}$ mice were euthanized by CO2 inhalation followed by cervical dislocation. L4-L6 lumbar dorsal root ganglia and spinal cord were dissected in PBS, and fixed for 15 and 30 min, respectively, in 4% paraformaldehyde (PFA) at room temperature. Tissues were rinsed twice in PBS before immersion overnight in 30% sucrose/PBS at 4°C. In situ hybridization was performed with standard procedures on transverse sections of 14 µm as described previsously10. Image acquisition was done using a Hamamatsu NanoZoomer using NDP view software. Analysis of *in situ* hybridization images : Slides containing DRG sections (12 µm) were originally imaged for endogenous mCherry fluorescence and then submitted to *in situ* hybridization using *Flt3* probes designed to target *Flt3* mRNA (*13*). After proper enzymatic staining, slides were re-imaged in bright field. Imaged were then processed as two separate channels, one for mCherry and one for Flt3 mRNA and superimposed using Photoshop software. Distortion was adjusted as needed to reach perfect matching between the two channels as hybridization treatment sometimes resulted in slight modification of overall DRG morphology. Finally, neurons positive for each or both channels were manually counted.

**Skin nerve preparation and ex vivo electrophysiological recordings**

The in vitro glabrous skin-nerve preparation used was adapted from Zimmermann *et al.* (*30*). Briefly, adult male mice were euthanatized using an overdose of isoflurane (Piramal, UK) followed by a method of confirmation of euthanasia. The saphenous nerve and the skin of the hind limb were carefully dissected and placed in a custom-designed organ chamber (INT-CFMN, Aix-Marseille University, Marseille, France) containing warm oxygenated Synthetic Interstitial Fluid (SIF buffer; 30.5°C). The SIF buffer had the following composition (in mM): 120 NaCl, 3.5 KCl, 0.7 MgSO4, 1.7 NaH2PO4, 5 Na2HCO3, 2 CaCl2, 9.5 Na-Gluconate, 5.5 glucose, 7.5 sucrose, and 10 HEPES. The pH was set to 7.4 and the osmolarity was maintained at 300 mOsm/L. The skin was placed with the corium side up in the organ bath and the saphenous nerve was placed in an adjacent recording chamber filled with mineral oil. The skin was continuously superfused with oxygenated SIF buffer at a temperature of 30.5°C controlled with a CL-100 temperature controller (Warner Instrument, Harvard apparatus, USA). The saphenous nerve was kept in a recording chamber filled with mineral oil, gently teased, and groups of nerve fibers were placed on the gold recording electrode in order to isolate single-unit activity. Then, extracellular action potentials from single nerve fibers were recorded with an AC differential amplifier DAM 80 (WPI) and converted digitally using the CED Spike2 system (sampling rate of 20 kHz; Cambridge Electronic Design, UK). Spikes were discriminated off-line with the Spike2 software (Cambridge Electronic Design Limited, UK) and analyzed individually to avoid false positives. Mechanical-threshold of single saphenous nerve fiber was obtained by probing the skin flap with von Frey hairs (Friedrich-Alexander University, Erlanger, Germany). Nociceptors were probed for their high threshold (> 16 mN) and tonic discharges during long lasting mechanical stimulation. Low-threshold mechanoreceptors (potential non nociceptors) were defined as having mechanical threshold below 5.7 mN and rapidly or slow adapting discharges. Once a fiber has been characterized, a single 10 s-supraliminar mechanical stimulation of the receptive field was applied using a von Frey filament mounted on the arm of a computer-controlled piezoelectric stepper (PI E-861 Nexact® controller; Germany). The post-discharge (or afterdischarge) was defined as a prolonged activity occurring within a 5 min-period after the end of the mechanical stimulus. Ratio of firing rates determined 5 min after the stimulus (post-discharge) and 5 min before the stimulus was used to analyze the post-discharge.

#### Slices preparation, electrophysiological recordings and stimulation

Adult male mice (3 to 6 weeks) were euthanatized with an intraperitoneal (i.p.) overdose of chloral hydrate (7 %). After a laminectomy, the thoraco-lumbar (T11-L3) spinal cord were removed and placed into an ice-cold sucrose-based saline solution containing the following (in mM): 2 KCl, 0.5 CaCl2, 7 MgCl2, 1.15 NaH2PO4, 26 NaHCO3, 11 glucose, and 205 sucrose bubbled with 95% O2, 5% CO2. Transverse slices (550 μm thick) with attached dorsal roots obtained using a vibratome (VT1200 S, Leica Microsystèmes SAS, France) were incubated at 37 °C in artificial cerebro-spinal fluid solution containing (in mM): 130 NaCl, 3 KCl, 2.5 CaCl2, 1.3 MgSO4, 0.6 NaH2PO4, 25 NaHCO3, 10 glucose (pH 7.4) and bubbled with 95 % O2 and 5 % CO2. After a 40 min recovery period, electrophysiological recordings were performed. Substantia gelatinosa neurons were visualized in spinal cord slices using an upright microscope fitted with fluorescence optics (AxioExaminer, Carl Zeiss, Germany) and linked to a digital camera QImaging Exi Aqua (Czech Republic). The lamina II was divided into outer (IIo) and inner (IIi) equal parts from the dorsal to ventral boundaries. Recorded lamina I and IIo interneurons were visualized using a 63x water immersion objective lens with combined infrared and differential interference. Patch pipettes (6-8 MΩ resistance) made from borosilicate glass (1.5 mm O.D; PG150T-15; Harvard Apparatus, UK) were filled with an internal solution containing (in mM): 127 K-gluconate, 4 NaCl, 2 Mg Cl2, 10 HEPES, 0.5 EGTA, 2.5 Na2ATP, 0.5 Na2GTP, 0.05% neurobiotin (Vector Laboratories), 0.01% dextran tetramethylrhodamine (10 000MW, fluoro-ruby, Life technologies), pH 7.4 and osmolarity 290-300mOsm. Whole-cell patch-clamp recordings were performed using Clampex 10 software connected to a Multiclamp 700B amplifier via a Digidata 1440A digitizer (Molecular Devices, Sunnyvale, CA, USA). The sampling rate was 10 kHz, and the data were filtered at 3 kHz. Series resistance was monitored throughout the experiments and data were discarded if series resistance varied more than ± 20 MΩ. In voltage-clamp mode, neurons were clamped at –65mV and evoked-postsynaptic currents (eEPSCs) were obtained by stimulating (5 pulses, 0.1ms, 0.05Hz, 500μA) the dorsal root attached with a suction electrode. Monosynaptic eEPSC were determined based on three criteria: constant short latency, a smooth waveform with a jitter up to 2ms and consistent responses without failure to repeated stimuli (*31*).

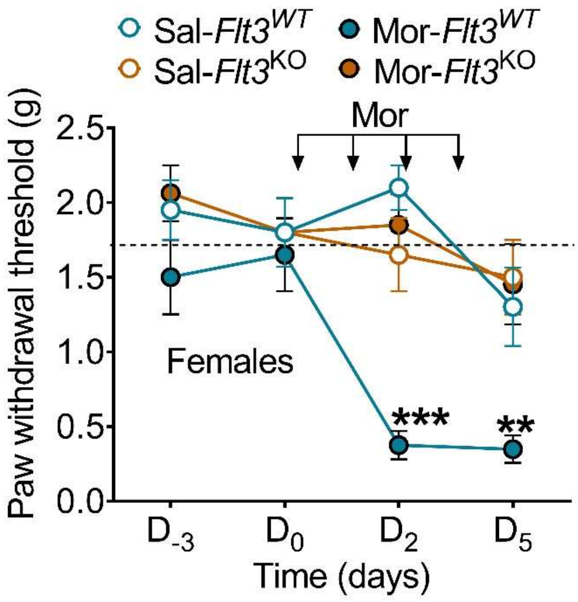


**Fig. S1: *Flt3* deletion prevents OIH in female mice.** Baseline nociceptive hypersensitivity of *Flt3*$^{WT}$ and *Flt3*$^{KO}$ female mice (n = 8 mice/group) measured by the von Frey « up and down » method after chronic morphine treatment (10mg/kg; subcutaneous, twice a day for 4 days). Data are shown as mean ± SEM. ***P < 0.001. Statistical analyses included two-way repeated ANOVA (**A**) followed by Bonferroni's test for multiple comparisons. Mor, morphine; Sal, saline.

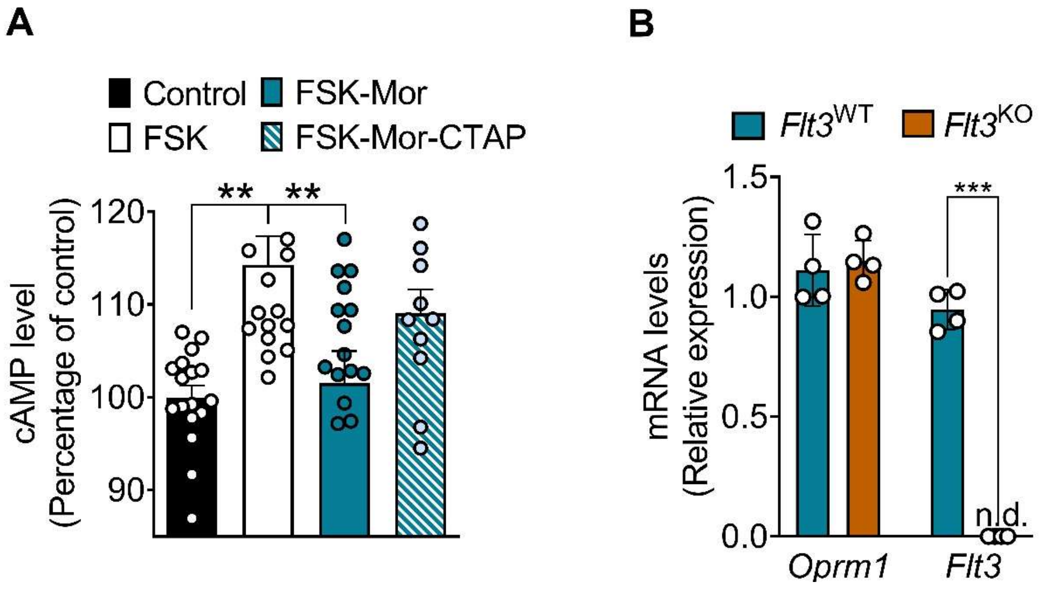


**Fig. S2: Effects of acute morphine on MOR-induced cAMP production in cultured sensory neurons and of Flt3 deletion on MOR gene expression. (A)** cAMP responses to forskolin in primary cultures of adult DRG neurons from $Flt3^{WT}$ in control, morphine and morphine + CTAP conditions (n = 4)**. (B)** Relative quantification of *Oprm1* and *Flt3* expression in DRG sensory neurons in *Flt3*$^{WT}$ and *Flt3*$^{KO}$ animals (n = 4 mice/group).. Data are shown as mean ± SEM. **P < 0.01, ***P < 0.001. Statistical analyses included one-way (**A**) or two-way (B) ANOVA followed by Bonferroni's test for multiple comparisons. FSK, forskolin; n.d., not detectable; Mor, morphine.

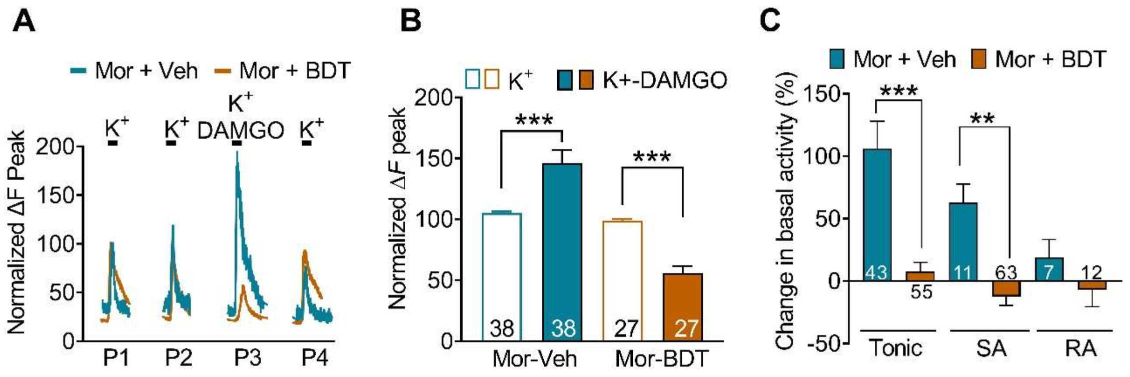


**Fig. S3: BDT001 inhibits MOR-induced hyperexcitability in mice. (A)** Traces of [Ca2+]i responses to repeated bath applications of high K+ (50 mM) alone or combined with DAMGO (1 µM), a MOR selective agonist, in chronic morphine-treated cultured DRG neurons (morphine at 10 µM for 3-4 DIV) with or without BDT001. (**B**) Results are expressed as response amplitudes normalized to HiK+ alone (ΔF peak) ((n= 4 mice). (**C**) *Ex vivo* extracellular recordings of mechanically evoked-activity on fibers from saphenous-nerve preparations from chronic morphine treated mice (10 mg/kg, subcutaneous twice a day for 4 days) with or without BDT001 (5 mg/kg, intraperitoneal, before and after a 10 s-mechanical stimulus applied on the skin (n = 3-7 mice/group). Data are represented as normalized firing of cutaneous mechanoreceptors. Data are shown as mean ± SEM. **P < 0.01, ***P < 0.001. Statistical analyses included mixed-effect model (REML) test followed by Šidák's for multiple comparisons (**B**) and one-way ANOVA followed by Bonferroni's test for multiple comparisons (**C**). BDT, BDT001; Mor, morphine; Veh, vehicle.

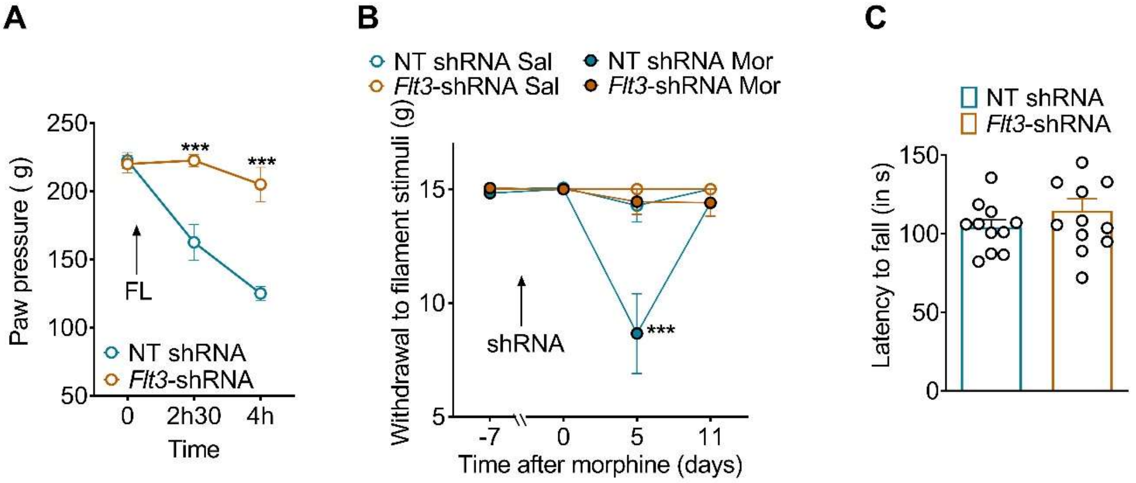


**Fig. S4: *Flt3*-targeted shRNA prevents FL (A) or morphine (B)-induced mechanical hyperalgesia and has no effect on motor function in rats.** Baseline mechanical hypersensitivity induced by intrathecal FL (**A**) or baseline mechanical allodynia induced by chronic morphine (**B**) in AAV9 non target-shRNA treated rats but not in AAV9 Flt3-shRNA treated rats (n = 6/group). (**C**) Motor analysis using the rotarod test in AAV9 non-target shRNA and AAV9 Flt3-shRNA treated rats. Data are shown as mean ± SEM. ***P < 0.001. Statistical analyses included two-way repeated ANOVA followed by Bonferroni's test for multiple comparisons (**A, B**) and Mann-Whitney test (**C**). Mor, morphine; NT, non-targeted;

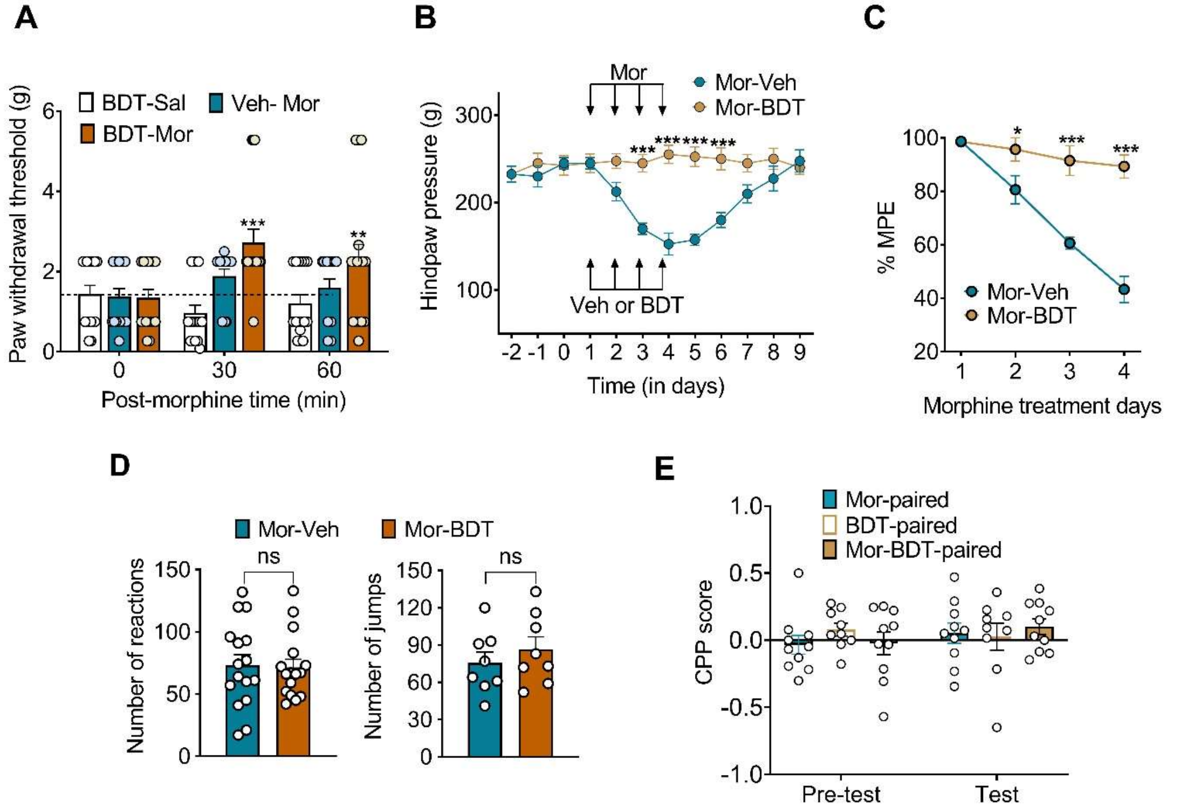


**Fig. S5: BDT001 potentiates morphine analgesia and prevents MIH and MIT without worsening morphine adverse effects**. (**A**) The analgesic effects of subcutaneous (s.c.) morphine (5 mg/kg) with or without intraperitoneal (i.p.) BDT001 (BDT, 5 mg/kg) were evaluated using von Frey "up and Down" method in mice (n=8/group). (**B**) Mechanical sensory sensitivity of rats receiving i.p. vehicle or BDT001 (5 mg/kg) 90 min before s.c. chronic morphine treatment (2mg/kg, twice a day for 4 days) (n=6 rats/group). (**C**) Measurement of morphine antinociception (percentage of MPE) during s.c. chronic morphine treatment (n=6 rats/group). (**D**) BDT001 (5 mg/kg) did not enhance naloxone-precipitated withdrawal signs in chronic morphine mice. (**E**) BDT001 (5 mg/kg) had no motivational effects and did not potentiate the motivational effects of a low, inactive dose of morphine (Mor 1.5 mg/kg) as evaluated with the conditioned place preference. Data are shown as mean ± SEM (n = 10-12 mice/group). **P* < 0.05, ***P* < 0.01, ****P* < 0.001 vs. morphine alone. Statistical analyses included two-way repeated ANOVA followed by Bonferroni's test for multiple comparisons (**A-C**), Mann-Whitney test (**D**), and two-way repeated ANOVA (**E**). BDT, BDT001; Mor, morphine; Sal, saline; Veh, vehicle.

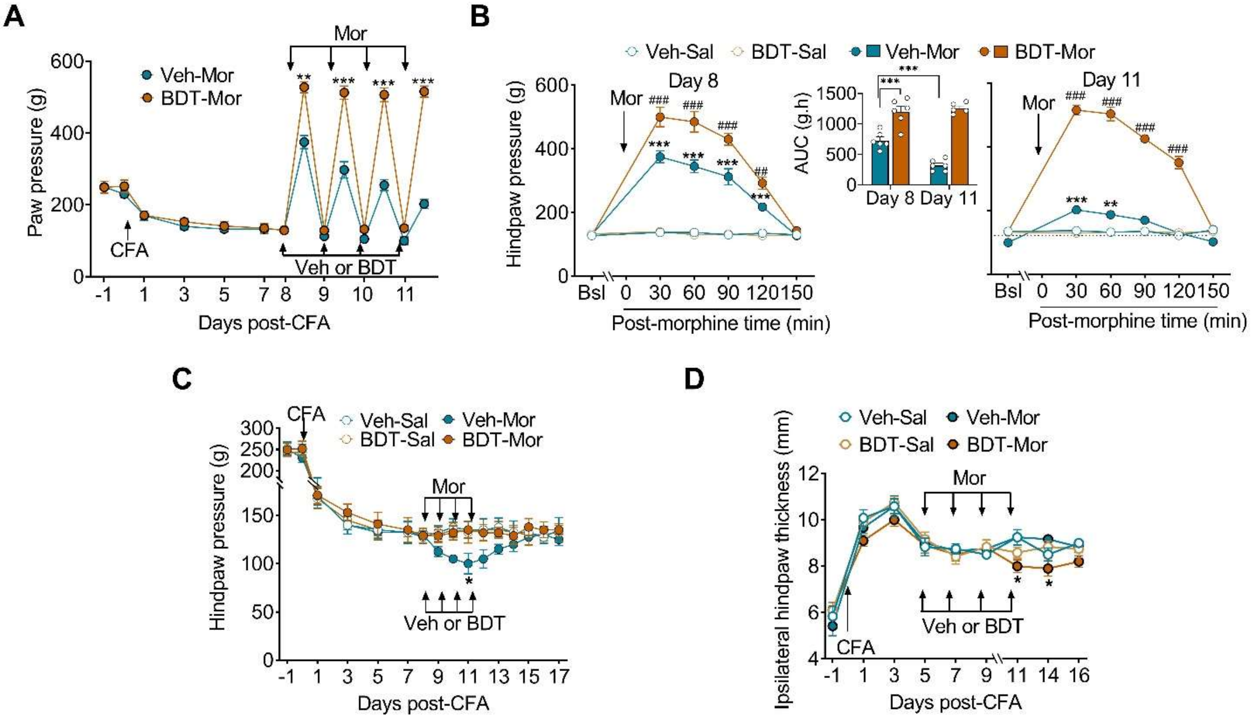


**Fig. S6: Preventive effects of BDT001 on MIH and MIT in a model of chronic inflammatory pain.** Rats received one injection of CFA in the hindpaw and treated with saline (Sal) or morphine (Mor, 5 mg/kg twice a day) and vehicle (Veh) or BDT001 (BDT, 5 mg/kg once a day) was initiated at 8 days after CFA ($D_8$) and repeated for 4 days. (**A**), Morphine analgesia progressively decreased, as morphine-induced increase in mechanical pain thresholds vanished, as shown by the maximal morphine effect at 30 min. BDT001-treated animals showed the maintenance of morphine analgesia (n = 6 rats/group). (**B**) The time-courses of morphine analgesic effect 8 and 11 days after CFA, with the area under the curves of the time-courses shown in the inset. MIT was completely prevented by BDT001. (**C**) When assessed before the first daily morphine administration, MIH progressively emerged with morphine treatment, as shown by the diminished mechanical pain threshold, both effects were completely prevented by BDT001. (**D**) BDT001 effects on morphine analgesia occurred without any sign of inflammation, as shown by the limited changes on paw thickness. Data are shown as mean ± SEM (5-6 rats/group). * *P < 0.05*, ***P* < 0.01, ****P* < 0.001 vs. Veh-Veh alone; ## *P* < 0.01, ### *P* < 0.001 vs. Veh-Mor. Statistical analyses included two-way ANOVA followed by Bonferroni's for multiple comparison test. ). BDT, BDT001; Mor, morphine; Sal, saline; Veh, vehicle.
